# First identification and investigation of piRNAs in the larval guts of Asian honey bee, *Apis cerana*

**DOI:** 10.1101/2022.10.30.514393

**Authors:** Qi Long, Ming-Hui Sun, Xiao-Xue Fan, Zong-Bing Cai, Kai-Yao Zhang, Si-Yi Wang, Jia-Xin Zhang, Xiao-Yu Gu, Yu-Xuan Song, Da-Fu Chen, Zhong-Min Fu, Rui Guo, Qing-Sheng Niu

## Abstract

Piwi-interacting RNAs (piRNAs), a kind of small non-coding RNAs (ncRNAs), play pivotal parts in maintaining the genomic stability and modulating biological processes such as growth and development via regulation of gene expression. However, piRNAs in Asian honey bee (*Apis cerana*) is still largely unknown at present. In this current work, on basis of previously gained high-quality small RNA-seq datasets, piRNAs in the larval guts of *Apis cerana cerana*, the nominate species of *A. cerana*, was for the first time identified, followed by in-depth investigation of the regulatory roles of differentially expressed piRNAs (DEpiRNAs) in the developmental process of the *A. c. cerana*. Here, a total of 621 piRNAs were identified in the *A. c. cerana* larval guts, among which 499 piRNAs were shared by 4- (Ac4 group), 5- (Ac5 group), and 6-day-old (Ac6 group) larval guts, while the numbers of unique ones were 79, 37, and 11, respectively. piRNAs each group were ranged from 24 nt to 33 nt in length, and the first base of piRNAs had a cytosine (C) bias. Additionally, five up-regulated and five down-regulated piRNAs were identified in the Ac4 vs. Ac5 comparison group, 9 of which could target 9, 011 mRNAs; these targets were involved in 41 GO terms and 137 pathways. Comparatively, 22 up-regulated piRNAs were detected in the Ac5 vs. Ac6 comparison group, 21 of which could target 28, 969 mRNAs; these targets were engaged in 46 functional terms and 164 pathways. The results suggested the overall alteration of expression pattern of piRNAs during the developmental process of *A. c. cerana* larvae. Regulatory network analysis showed that piR-bmo-748815 and piR-bmo-512574 in the Ac4 vs. Ac5 comparison group as well as piR-bmo-716466 and piR-bmo-828146 in the Ac5 vs. Ac6 comparison group linked to the highest number of targets. Further investigation indicated that targets of DEpiRNAs in the above-mentioned two comparison groups could be annotated to several growth and development-associated pathways, such as Jak/STAT, TGF-β, and Wnt signaling pathways, indicating the involvement of DEpiRNAs in modulating larval gut development via these crucial pathways. Moreover, the expression trends of six randomly selected DEpiRNAs were verified using a combination of stem-loop RT-PCR and RT-qPCR. These results not only provide a novel insight into the development of the *A. c. cerana* larval guts, but also lay a foundation for uncovering the epigenetic mechanism underlying the larval gut development.

## 1. Introduction

Piwi-interacting RNAs (piRNAs), a kind of small non-coding RNA (ncRNA) with a length of 23-31 nucleotides (nt), are abundant in germ cells and reproductive tissues [1]. PiRNAs are generated from single-stranded precursor transcripts independently of Dicer and possess 2’-O-methylation at their 3’ end. PiRNAs were firstly identified in *Drosophila melanogaster*, and then in *mice* [1], *Droshplia* [2]and *Caenorhabditis elegans* [3]. In insects, limited work was mainly relevant to a few model species such as *Droshplia* [2], silkworms [4] and flies [5]. However, study on other insects including honey bee is still at the preliminary stage. A major function of piRNAs is to maintain the genome stability via suppressing the activity of transposons [6]. In recent years, with the development of related knowledge and technology, piRNAs were suggested to modulate gene expression through targeting mRNAs, similar to the action manner of miRNAs, another kind of small RNAs [7]. For example, Manage et al. found that SIMR-1 interacted PRG-1 in the piRNA pathway to promote the expression of downstream target genes by the mutator complex [8]; Shen et al. discovered that piRNAs in *C. elegans* could target all mRNAs in germline following miRNA-like pairing rules [9].

*Apis cerana* is wildly reared in China and many other Asian countries, playing an essential role in ecology and economy through pollination for wild flowers and crops as well as production of api-products [10].Previously, our group conducted a series of studies on ncRNAs regulating larval development and immune defense of *A. c. cerana*, the nominate subspecies of *A. cerana*, e.g., Feng et al. performed transcriptome-wide identification of miRNAs in the *A. c. cerana* larval guts followed by validation of the expression and sequence of 6 miRNAs [11]; Du et al. deciphered the differential expression profile of lncRNAs in the *A. c. cerana* larval gut in response to *Ascosphaera apis* infection [12]; Chen et al. analyzed the expression patterns of circRNAs in the larval gut of *A. c. cerana* responding to *A. apis* invasion and unraveled the putative function of differentially expressed circRNAs (DEcircRNAs) in host immune response [13]. Compared with *Drosophila* and *Aedes albopictus*, study on honey bee piRNAs is still limited. Liao et al. identified two PIWI genes, *Am-aub* and *Am-ago3*, for the first time in worker, drone, and queen bees of *A. mellifera*, and the expression of these two genes showed a significant sex bias, confirming that piRNA are truly expressed in western honey bee, indicating that piRNA may have a potential association with honey bee reproduction under the influence of nutritional factors as well as bee sex determination [14]; Based on RT-qPCR detection, Wang et al. identified piRNA clusters content among drone, worker and queen, found that the expression level of piRNAs in reproductive individuals of *A. mellifera* was greater than that in sterile workers, suggesting the reproductive bias of piRNA expression[15]; Watson et al. further surveyed the expression level of piRNAs in various *A. mellifera* reproductive tissues such as ovaries, spermatheca, semen, fertilised eggs, unfertilised eggs, and testes, and observed that [16]. Recently, on basis of transcriptome data and bioinformatics, our team identified 843 piRNAs in of the larval guts of *Apis mellifera ligustica* and uncovered the potential roles of DEpiRNAs during the developmental process of larval guts [17]. However, little progress piRNAs in *A. cerana*, the sister species of *A. mellifera* has been made until now.

To decipher the differential expression pattern of piRNA during the developmental process of *A. c. cerana* larval guts and the regulatory roles of DEpiRNAs, based on our previously gained high-quality small RNA-seq datasets, identification and structural analysis of piRNAs in *A. c. cerana* larval guts were conducted. In this current work, followed by investigation of DEpiRNAs and discussion of their potential roles in regulation of larval gut development. To our knowledge, this is the first documentation of *A. cerana* piRNAs. Findings in the present study will enrich the reservoir of *A. cerana* piRNAs and deepen our understanding of ncRNA-modulated development of larval guts.

## 2. Materials and Methods

### 2.1 Transcriptome data source

*A. c. cerana* larvae used in this study were obtained from colonies reared in the apiary at the College of Animal Sciences (College of Bee Science), Fujian Agriculture and Forestry University, Fuzhou city, China. In a previous work, 4-, 5-, and 6-days-old (Ac4, Ac5, and Ac6 groups) larval gut tissues were respectively prepared and subjected to RNA isolation, cDNA library construction, and deep sequencing using sRNA-seq technology, there were three biological replicas of each group, and each group contained three larval guts [11]. Totally, 11, 273, 306; 11, 349, 964 and 11, 122, 092 raw reads were generated from Ac4, Ac5 and Ac6 groups, and 9, 394, 648; 36, 9, 402, 531 and 9, 394, 648 clean tags were respectively gained after quality control [11]. Raw data generated from sRNA-seq were deposited in the NCBI SRA database and linked to the BioProject number: No: PRJNA395108.

### 2.2 Identification and analysis of piRNAs

Following the previously described method by Xu et al.[17], (1) the clean reads from each group were aligned to the *A. cerana* reference genome (Assembly ACSNU-2.0) to obtain mapped reads, and then other types of sRNAs were filtered; (2) small ncRNAs, such as rRNA, scRNA, snoRNA, snRNA, and tRNA, were filtered out by mapping the remaining clean reads to GeneBank and Rfam (11.0) databases; (3) miRNAs in the remaining clean reads were further filtered out; (4) sRNAs between 24 nt and 33 nt in length were retained according to the length characteristics of piRNA, and finally, only sRNAs mapped to a unique position were retained as candidate piRNAs.

The expression of each piRNA was normalized using the TPM method (TPM=T * 106/N, T stands for clean reads of piRNA and N stands for clean reads of total sRNAs). Subsequently, the structural features of piRNAs including length distribution and first base bias were analyzed based on the prediction result. Upset plot and venn analysis of piRNA expression level were visualized using omicshare platform (https://www.omicshare.com/tools/)

### 2.3 Investigation and target prediction of DEpiRNAs

Following to the criteria of |log2 fold change| ≥ 1 and *P* ≤ 0.05, DEpiRNAs in Ac4 vs. Ac5 and A5 vs. Ac6 comparison groups were screened out. Target mRNAs of DEpiRNA were then predicted using Targetfinder software [18] with default parameters. Next, using the BLAST tool, target mRNAs were respectively aligned to GO (https://www.geneontology.org) and KEGG (https://www.genome.jp/kegg/) databases, to gain functional and pathway annotations.

### 2.4 Analysis of DEpiRNA-mRNA regulatory network

Based on the predicted targeting relationships, regulatory networks between DEpiRNA and target mRNAs were constructed following the thresholds of free energy < −20 kcal mol −1 and *P* < 0.05, followed by visualization using Cytoscape software. Further, on basis of the KEGG pathway annotations, the target mRNAs annotated in Wnt, TGF-β and Hippo signaling pathways were further surveyed to construct corresponding regulatory networks, which were then visualized with Cytoscape software [19].

### 2.5 Stem-loop RT-PCR of piRNAs

Total RNA from 4-, 5-, and 6-day-old *A. c. cerana* larval guts were extracted using a FastPure® Cell/Tissue Total RNA Isolation Kit V2 (Vazyme, Nanjing, China). The concentration and purity of RNA were checked with Nanodrop 2000 spectrophotometer (Thermo Fisher, Waltham, MA, USA). three DEpiRNAs (piR-bmo-1010100, piR-ame-1183555 and piR-bmo-202265,) from Ac4 vs. Ac5 comparison group were randomly selected for stem-loop RT-PCR validation, and three (piR-bmo-828146, piR-bmo-904144 and piR-ame-11093) from Ac5 vs. Ac6 comparison group. Specific stem-loop primers and forward primers (F) as well as universal reverse primers (R) were designed using DNAMAN software and then synthesized by Sangon Biotech (Sangon Biotech, Shanghai, China). According to the instructions of HiScript ® 1st Strand cDNA Synthesis Kit, cDNA was synthesized by reverse transcription using stem-loop primers and used as templates for PCR of DEpiRNA. Reverse transcription was performed using a mixture of specific loop primers and oligo (dT) primers, and the resulting cDNA was used as templates for PCR. The PCR system (20 μL) contained 1 μL of diluted cDNA, 10 μL of PCR mix (Vazyme, Nanjing, China), 1 μL of forward primers, 1 μL of reverse primers, and 7 μL of diethyl pyrocarbonate (DEPC) water. The PCR was conducted on a T100 thermocycler (Bio-Rad, Hercules, CA, USA) under the following conditions: pre-denaturation step at 95 w for 5 min; 40 amplification cycles of denaturation at 95 w for 10s, annealing at 55 w for 30 s, and elongation at 72 w for 1 min, followed by a final elongation step at 72 w for 10 min. The amplification products were detected on 1.8% agarose gel electrophoresis with Genecolor (Gene-Bio, Shenzhen, China) staining.

### 2.6 RT-qPCR detection of DEpiRNAs

The RT-qPCR was carried out following the protocol of SYBR Green Dye kit (Vazyme, Nanjing, China). The reaction system (20 μL) included 1 μL of cDNA, 1 μL of forward primers, 1 μL of reverse primers, 7 μL of DEPC water, and 10 μL of SYBR Green Dye. RT-qPCR was conducted on an Applied Biosystems QuantStudio 3 system (Thermo Fisher, Waltham, MA, USA) following the conditions: pre-denaturation step at 95 w for 5 min, 40 amplification cycles of denaturation at 95 w for 10 s, annealing at 60 w for 30 s, and elongation at 72 w for 15 s, followed by a final elongation step at 72 w for 10 min. The reaction was performed using an Applied Biosystems QuantStudio 3 Real-Time PCR System (Thermo Fisher, Waltham, MA, USA). snRNA U6 was selected as inner reference gene. All reactions were performed in triplicate. The relative expression of piRNA was calculated using the 2-ΔΔCt method [20]. Detailed information about primers used in this work is shown in **Table S1**.

### 2.7 Statistical analysis

Statistical analyses were conducted with SPSS software (IBM, Amunque, NY, USA) and GraphPad Prism 7.0 software (GraphPad, San Diego, CA, USA). Data were presented as mean ± standard deviation (SD). Statistics analysis was performed using Student’s t test. Significant (*P* < 0.05) GO terms and KEGG pathways were filtered by performing Fisher’s exact test with R software 3.3.1 [21-22].

## 3. Results

### 3.1 Identification, structural analysis, and validation of piRNAs in A. c. cerana larval guts

Totally, 621, 558, and 549 piRNAs were identified in Ac4, Ac5, and Ac6 groups, respectively. In addition, 499 piRNAs were shared by the aforementioned three groups, whereas the quantities of specific ones were 79, 37, and 11, respectively **(Figure 1)**.

**Figure 1.**
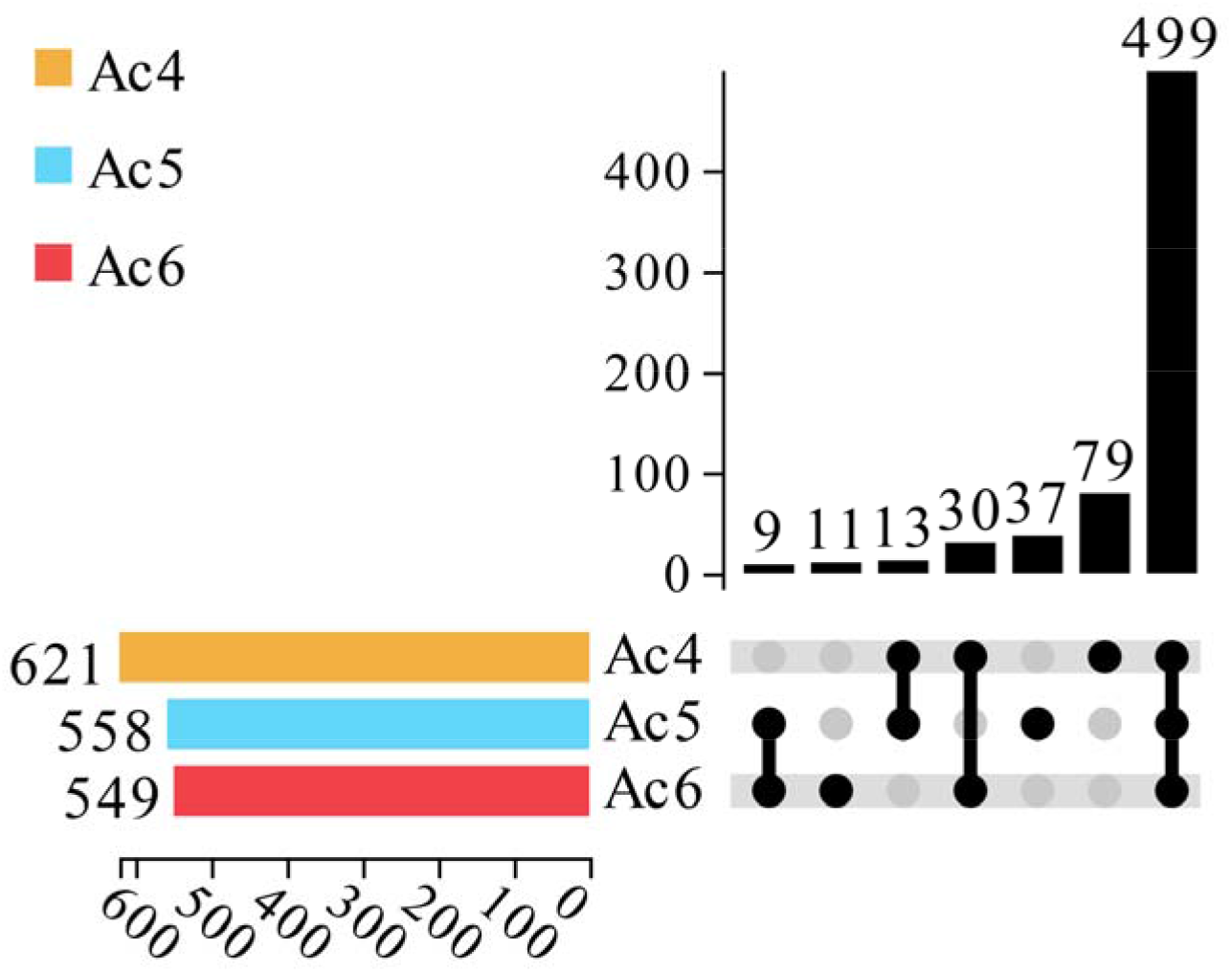
Upset plot of differential piRNA expression in *A. c. cerana* worker larval mid gut.

The lower left color column indicates the number of piRNA contained in different groups; the lower right node indicates the piRNA common to each group; the connection between each group indicates the common piRNA; the teamless node indicates the unique piRNA of this group; and the number of unique and common piRNA is presented above; they are arranged in ascending order from left to right.

Structural analysis indicated that the length distribution of the identified *A. c. cerana* piRNAs in the above-mentioned three groups were from 24 nt to 33 nt **(Figure 2A-C)**; additionally, the first base of piRNAs in 4, Ac5, and Ac6 groups had a C bias **(Figure 2E-F)**.

**Figure 2.**
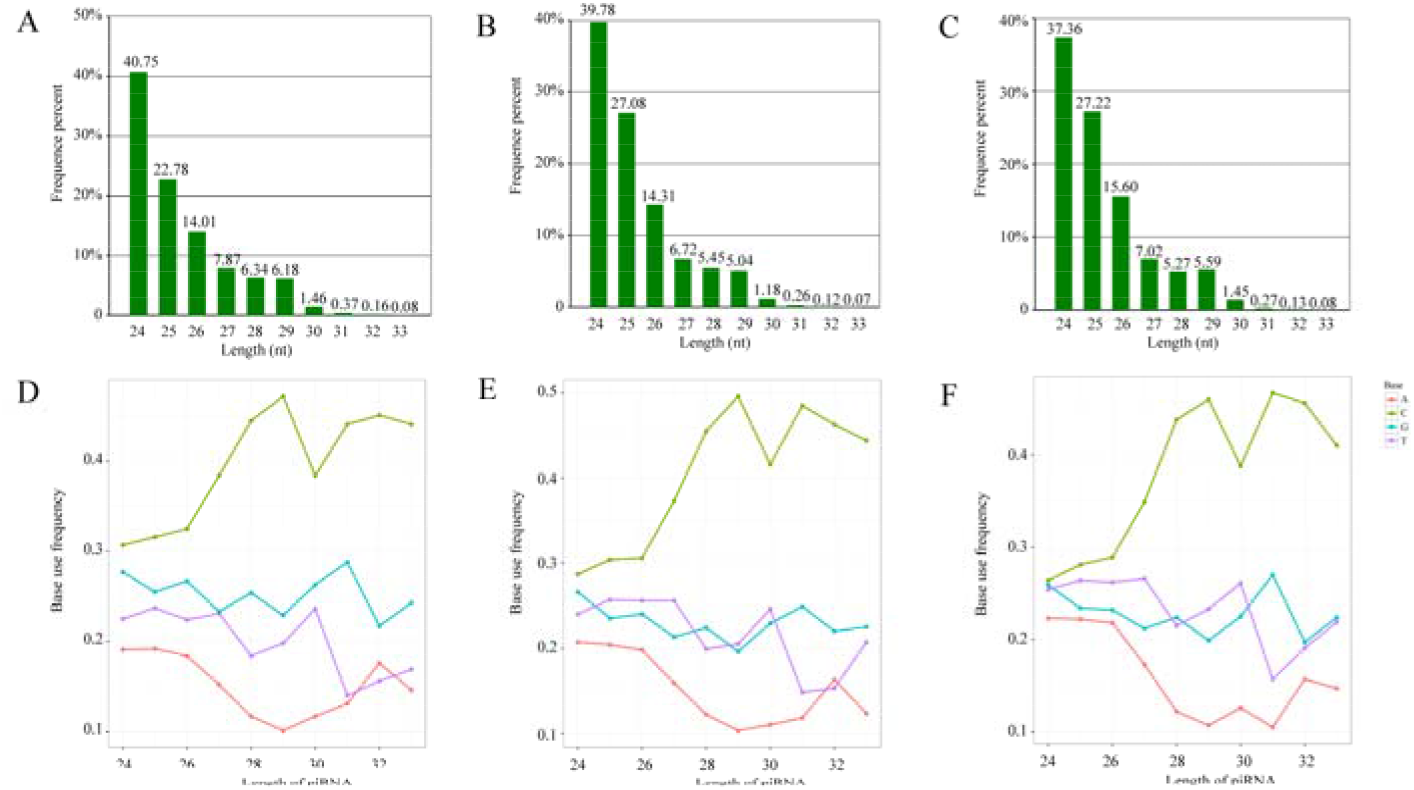
Length distribution and first base bias of *A. c. cerana* piRNAs. (A-C) Length distribution of piRNAs identified in Ac4, Ac5, and Ac6 groups. (D-F) first base bias of piRNAs identified in Ac4, Ac5, and Ac6 groups.

### 3.2 Differential expression pattern of piRNAs during the developmental process of A. c. cerana larval guts

Ten DEpiRNAs were discovered in the Ac4 vs. Ac5 comparison group, including six up-regulated and four down-regulated ones; among these, the most significantly up-regulated three piRNAs were piR-bmo-1183555 (log2FC= 1.816, *P*= 0.001), piR-bmo-458012 (log2FC= 1.469, *P*= 0.003), and piR-bmo-512574 (log2FC= 1.469, *P*= 0.003; while the three most significantly down-regulated ones were piR-bmo-1103846 (log2FC= −12.019, *P*= 5.32E-06), piR-bmo-762269 (log2FC= −1.251, *P*= 0.043), and piR-bmo-1010100 (log2FC= −1.228, *P*= 0.048) (Figure 3A). Comparatively, 22 up-regulated DEpiRNAs were characterized in the Ac5 vs. Ac6 comparison group, whereas no down-regulated one was detected; the most significantly up-regulated DEpiRNA was piR-bmo-750627 (log2FC=1.145, *P*= 0.002) followed by piR-bmo-11093 (log2FC= 1.666, *P*= 0.006) and piR-bmo-748816 (log2FC= 1.084, *P*= 0.006) **(Figure 3B)**. In addition, there were three up-regulated piRNAs shared by the two comparison groups mentioned above **(Figure 3C)**. Detailed information about piRNAs identified in this work were showed in **Table S2**.

**Figure 3.**
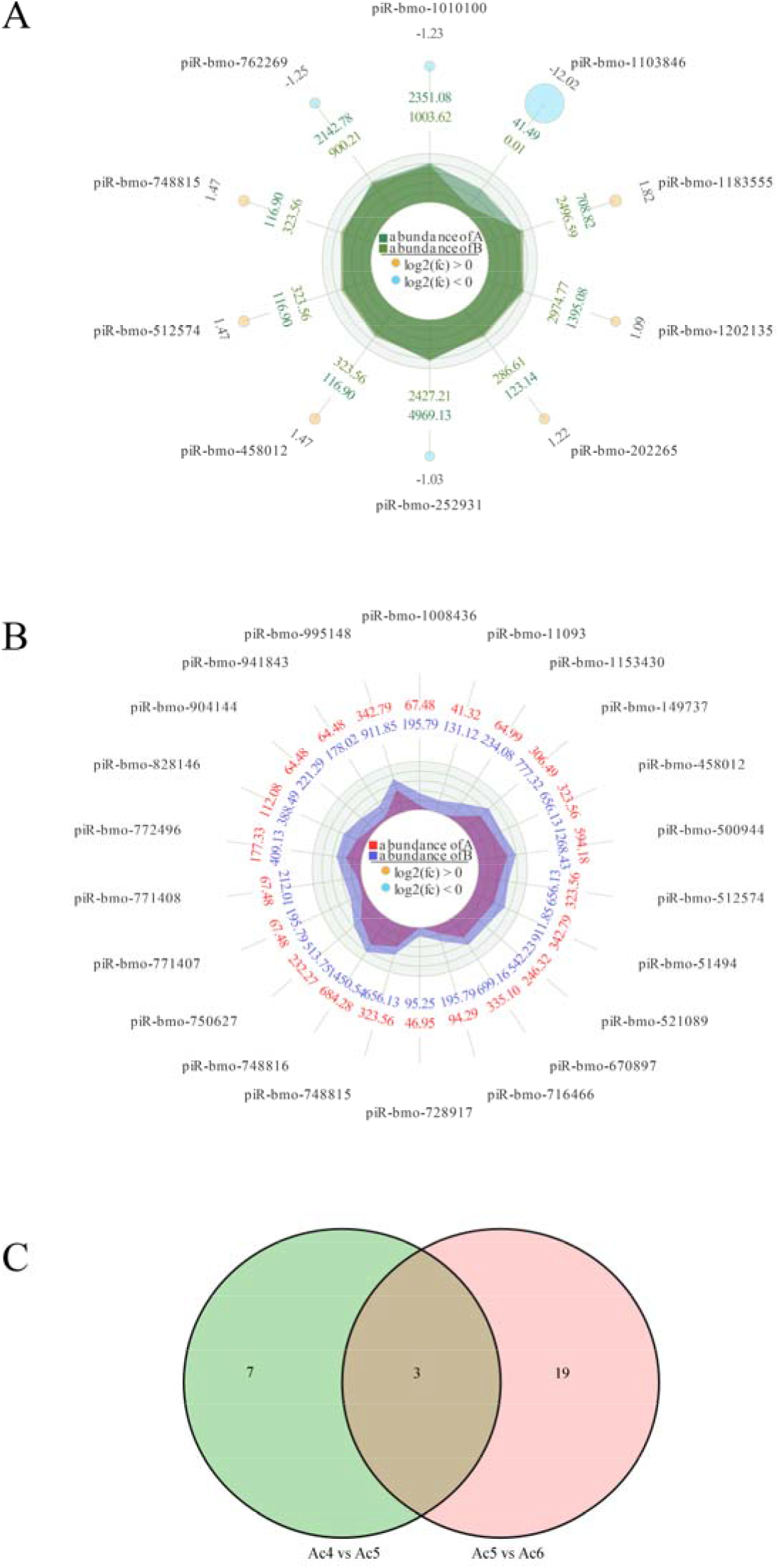
Radar maps and Venn diagram of DEpiRNAs. (A) Radar map of DEpiRNAs in Ac4 vs. Ac5 comparison groups. (B) Radar map of DEpiRNAs in Ac5 vs. Ac6 comparison groups. (C) Venn analysis of DEpiRNAs in two comparison groups.

### 3.3 Analysis and annotation of DEpiRNA-targeted genes

In the Ac4 vs. Ac5 comparison group, nine DEpiRNAs were found to target 9, 011 mRNAs, which could be annotated to 17 biological process-associated GO terms such as biological adhesion and biological regulations, ten molecular function-associated terms such as binding and transcription factor activity, protein binding, and 14 cellular component-associated terms such as membrane part and membrane **(Figure 4A)**. In comparison, 21 DEpiRNAs in the Ac5 vs. Ac6 comparison group were detected to target 28, 969 mRNAs, which could be annotated to 20 function term relative to biological process such as cellular process and localization, ten terms relevant to molecular function such as transporter activity and molecular function regulator, and 16 terms related to cellular component such as synapse part and. **(Figure 3B)**.

**Figure 4.**
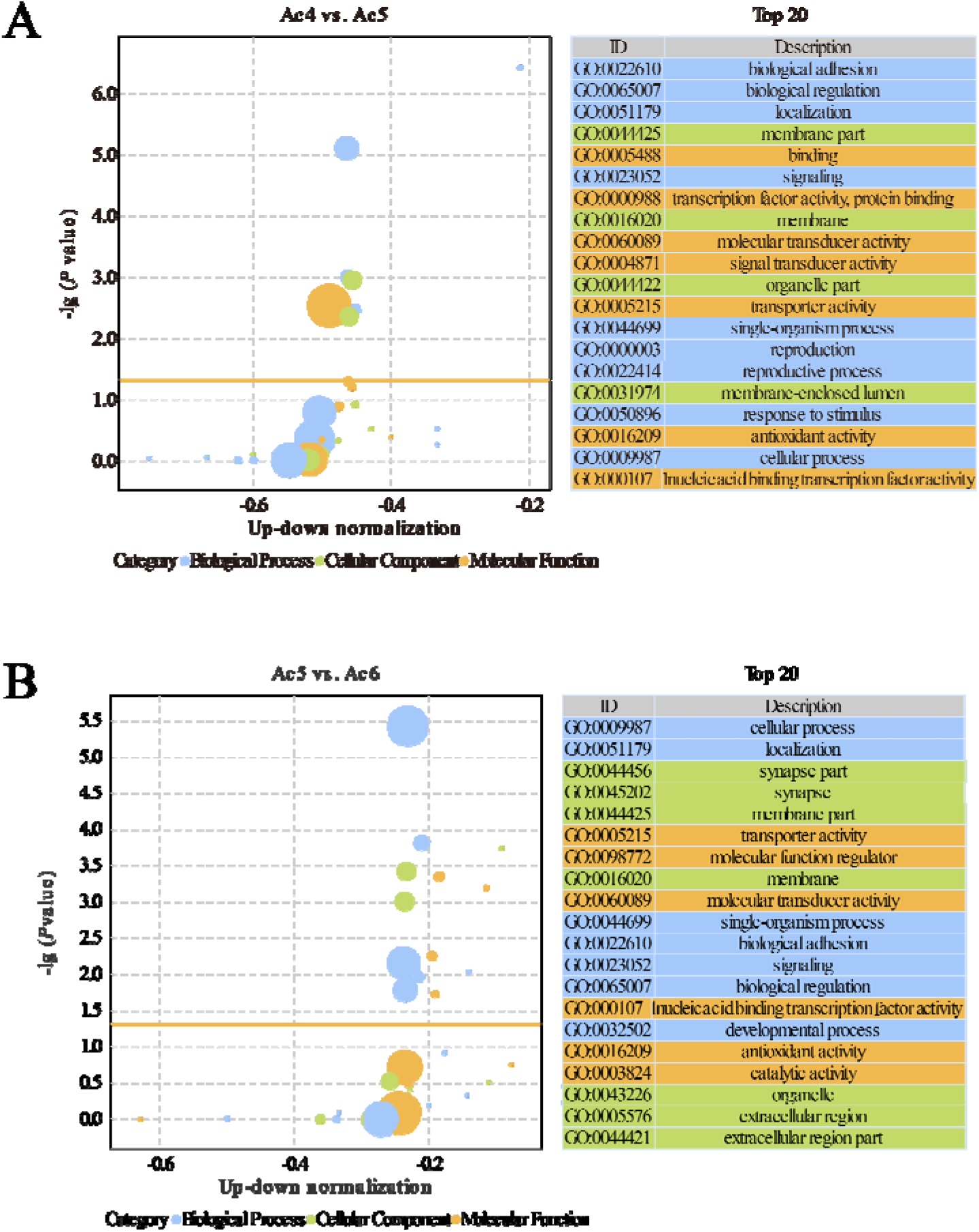
GO terms annotated by target mRNAs of DEpiRNAs. (A) Bubble diagram of DEpiRNA-targeted mRNAs in Ac4 vs. Ac5 comparison group; (B) Bubble diagram of DEpiRNA-targeted mRNAs in Ac5 vs. Ac6 comparison group.

Additionally, target genes of DEpiRNAs in Ac4 vs. Ac5 comparison group could be annotated to 137 KEGG pathways associated with environmental information processing, metabolism, organismal systems, genetic information processing, human diseases and cellular processes, such as endocytosis, fatty acid biosynthesis, and Wnt signaling pathway **(Figure 5A)**. Comparatively, DEpiRNA-targeted mRNAs in Ac5 vs. Ac6 comparison group could be annotated to 164 pathways including the AGE-RAGE signaling pathway in diabetic complications, ubiquitin mediated proteolysis, and TGF-beta signaling pathway **(Figure 5B)**.

**Figure 5.**
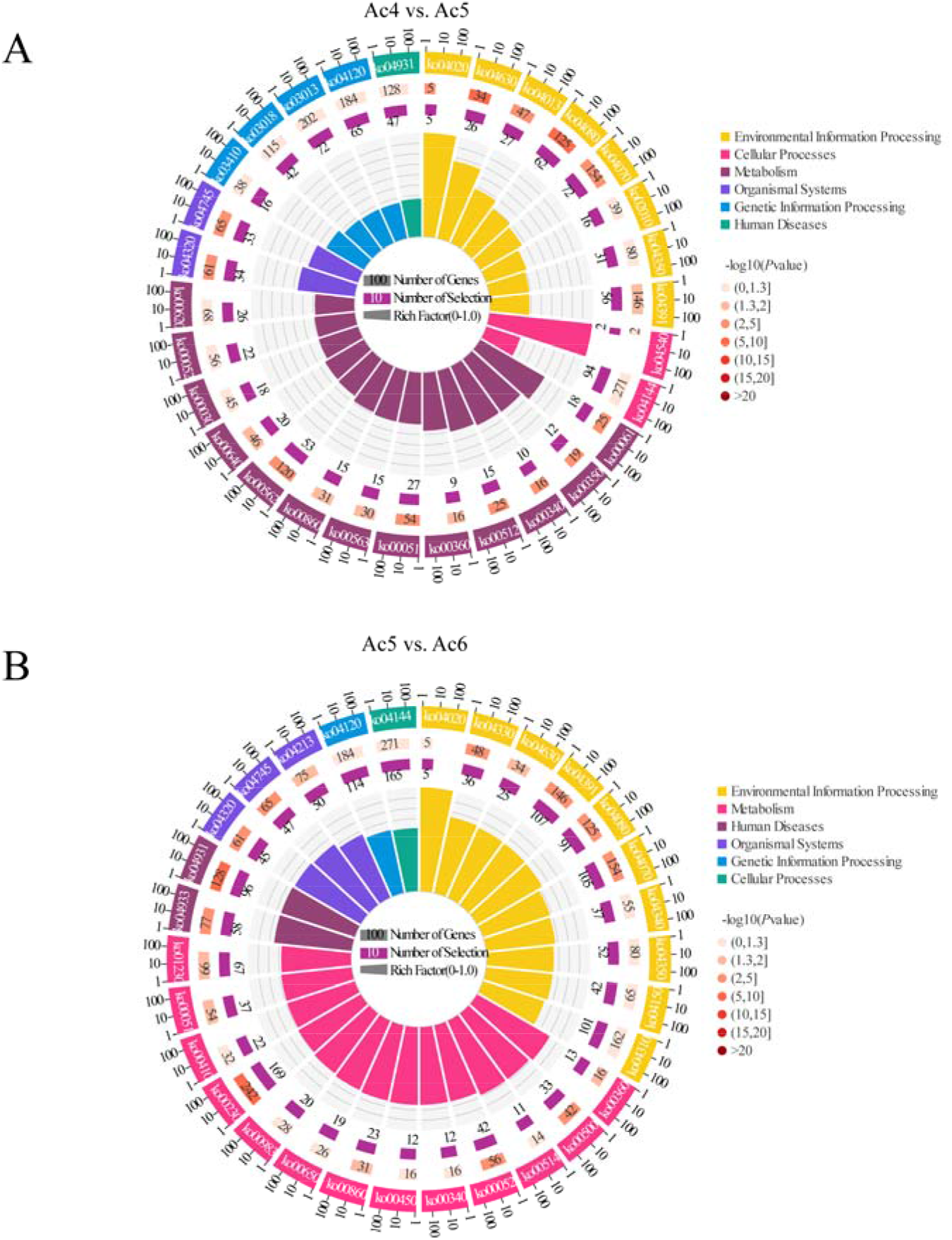
KEGG pathways annotated by target mRNAs of DEpiRNAs. (A) Pathways annotated by DEpiRNA-targeted mRNAs in Ac4 vs. Ac5 comparison group; (B) Pathways annotated by DEpiRNA-targeted mRNAs in Ac5 vs. Ac6 comparison group.

### 3.4 Regulatory network between DEpiRNAs and target genes

In the Ac4 vs. Ac5 comparison group, each DEpiRNA can target multiple mRNAs, with piR-bmo-748815 and piR-bmo-512574 binding to the highest number of targets (1, 701 and 1, 718). Similarly, each DEpiRNA in the Ac5 vs. Ac6 comparison group can target several mRNAs, with piR-bmo-716466 and piR-bmo-828146 linking to the highest number of targets (2, 089 and 2, 620).

As shown in Figure, there were complex regulatory relationships between DEpiRNAs and target mRNAs in the aforementioned two comparison groups. Further analysis demonstrated that 105 and 178 target mRNAs in the above-mentioned two comparison groups were involved in Wnt signaling pathway, Jak/STAT pathway and TGF-β signaling pathway **(Figure 6)**. Detailed information about DEpiRNAs and corresponding target mRNAs were presented in **Table S3**.

**Figure 6.**
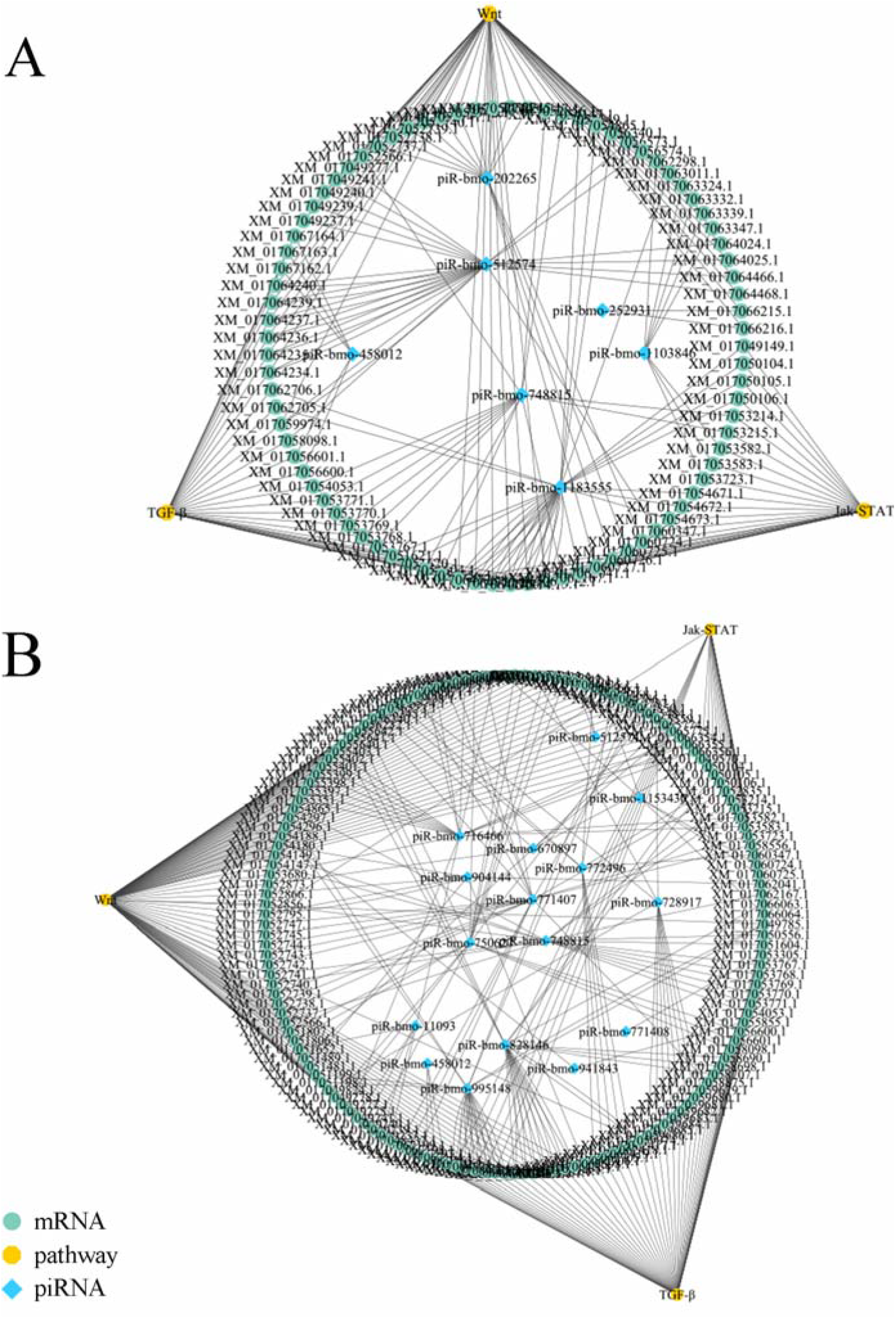
Regulatory network of *A. c. cerana* worker larvae. (A) Regulatory network of DEpiRNAs in Ac4 vs. Ac5; (B) Regulatory network of DEpiRNAs in Ac5 vs. Ac6

### 3.5 *Verification of DEpiRNAs* by *stem-loop RT-PCR and RT-qPCR*

Six randomly selected DEpiRNAs were subjected to stem-loop RT-PCR verification, the result was indicative of their expression in the larval gut **(Figure 7)**.

**Figure 7.**
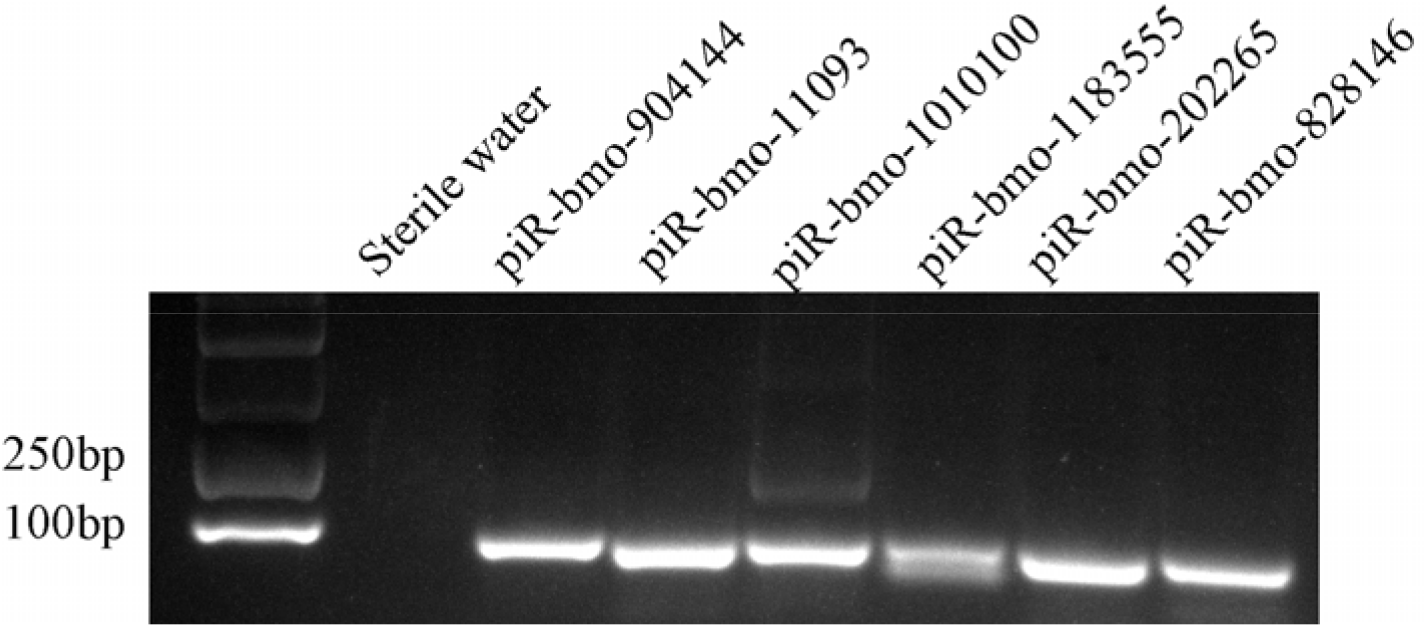
Stem-loop RT-PCR validation of six *A. c. cerana* DEpiRNAs.

**Figure 8.**
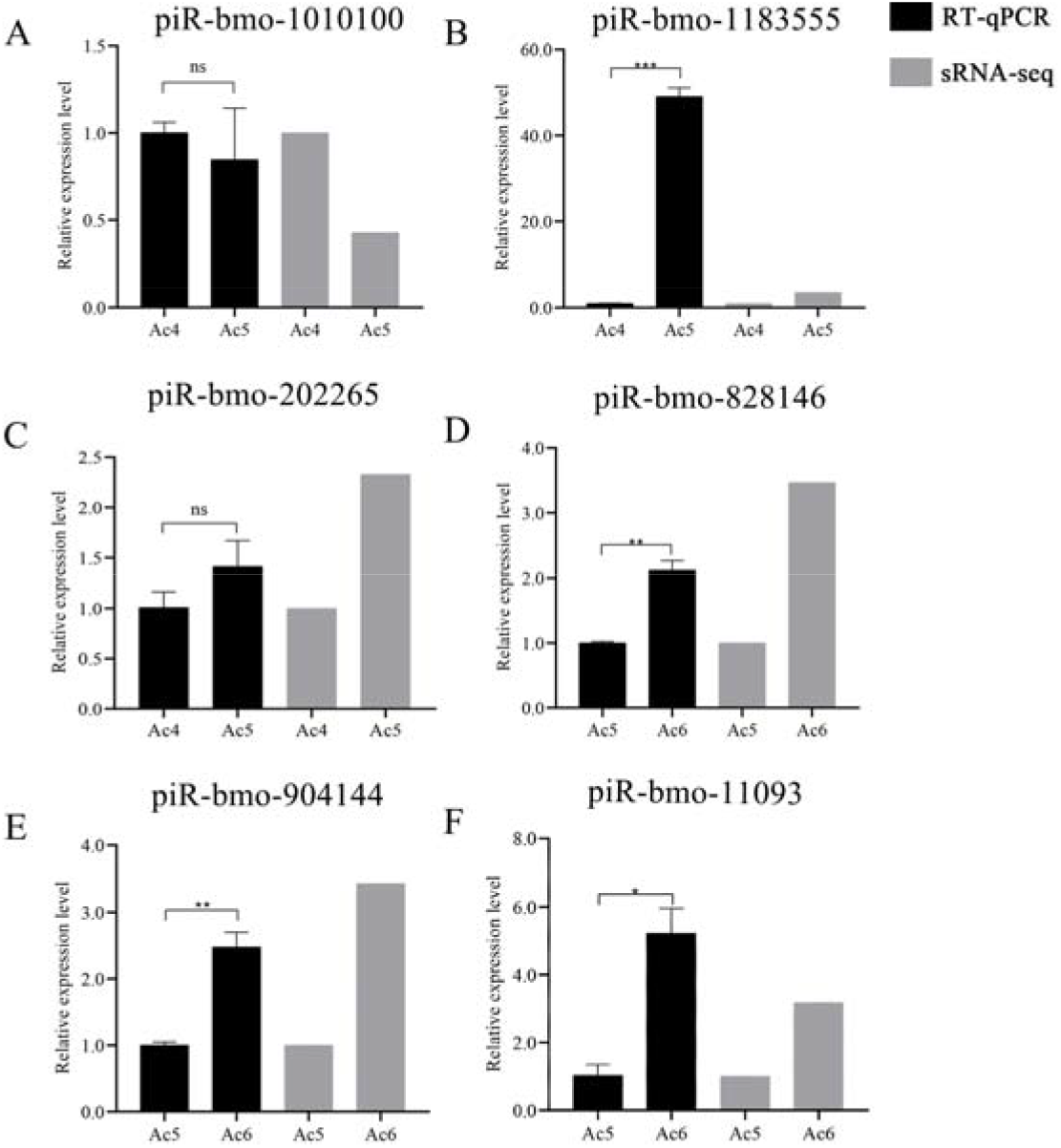
RT-qPCR verification of six *A. c. cerana* DEpiRNAs. (A-C) Relative expression levels of piRNAs from the Ac4 vs. Ac5 comparison group. (D-F) Relative expression level of piRNA from the Ac5 vs. Ac6 comparison group. ns indicates non-significant, * indicates *p* < 0.05, ** indicates *p* < 0.01 and *** indicates *p* < 0.001.

Further, RT-qPCR result suggested that the expression trends of these six DEpiRNAs were consistent with those in transcriptome datasets, confirming the authenticity and reliability of our sequencing data.

## 4. Discussion

Previous studies on honey bee piRNAs were mainly associated with *A. mellifera*. Here, on basis of our previously gained sRNA-seq data, we for the first time identified 621, 558, and 549 piRNAs in 4-, 5-, and 6-day-old *A. c. cerana* larval guts, respectively. After removing redundant ones, a total of 621 *A. c. cerana* piRNAs were obtained, offering valuable resource for further functional investigation and mechanism exploration in the future.Gonad tissues were considered to be the primary generative location of piRNAs. However, accumulating evidence indicated that piRNAs were also abundantly expressed in other organs and tissues of animals [23]. Morazzani et al. revealed that piRNAs were widespread and abundant in the mosquito soma by next-generation sequencing [24]. Feng et al. identified 3, 396 piRNA in the midgut tissues of *B. mori* using Illumina sequencing and bioinformatics [23]. We observed that the overall expression levels of the identified piRNAs in *A. c. cerana* larval gut displayed an up-regulation-down-regulation trend (Figure S1), this was indicative of the involvement of piRNAs in the development of *A. c. cerana* larval guts. Comparatively, although more piRNAs were identified in *A. m. ligustica*, the overall expression levels of piRNAs in the guts of *A. m. ligustica* 4-, 5-, and 6-day-old larvae was more consistent during development [17]. Together, these results demonstrated that piRNAs had different dynamics during the developmental process of larval guts of the above-mentioned two different bee species, implying different piRNA-regulated mechanisms underlying larval gut development.

An array of studies suggested that piRNAs can exert regulatory functions by targeting mRNAs via base pairing in various species, such as *Drosophila* [25], *C. elegans* [26] and mouse [27]. In silkworms, piRNAs were reported to be involved in sex determination through down-regulation of target genes [28]; down-regulation of target mRNAs by piRNA was observed during sex determination [28]. In addition, it’s proposed that piRNAs could also interact with targets without causing cleavage following perfect base-pairing [26]. A recent study in *C. elegans* defined the piRNA seed region from the 2nd to 7th nucleotide and observed that base pairing outside of the seed region contributed to piRNA target recognition [26]. Here, on basis of bioinformatics, nine and 21 piRNAs in Ac4 vs. Ac5 and Ac5 vs. Ac6 comparison groups were predicted to respectively target 9, 011 and 28, 969 mRNAs; among these, piR-bmo-748815 and piR-bmo-1183555 were found to bind to the highest number of targets, followed by piR-bmo-716466 and piR-bmo-828146 (Table S3), indicative of the highest connectivity of these DEpiRNAs, which deserved additional investigation.

Wnt signaling pathway plays essential functions in a number of biological processes, from embryogenesis and adult homeostasis to regulate cell proliferation, cell polarity, and the specification of cell fate [29]. In insects, Wnt signaling pathway was found to exert regulatory functions in developmental process, eg. Oberhofer et al. reported that Wnt signaling pathway activates germ layers related genes, including pair-rule, Tc-caudal and Tc-twist genes, further acts on growth zone metabolism and cell division, indicated that Wnt pathway is required for hindgut development [30]. Here, it’s detetcted that mRNAs relevant to Wnt signaling pathway were targeted by 6 DEpiRNAs (piR-bmo-1183555, piR-bmo-512574 and piR-bmo-202265, piR-bmo-748815, piR-bmo-1103846 and piR-bmo-252931) in the Ac4 vs. Ac5 comparison group and 21 DEpiRNA (piR-bmo-11093, piR-bmo-24995 and piR-bmo-748814, etc.) in the Ac5 vs. Ac6 comparison group, which suggested that these DEpiRNAs were potentially engaged in regulating the development of *A. c. cerana* larval guts via Wnt signaling pathway. More efforts are needed to decipher the underlying mechanism.

Signaling pathways are highly interconnected and extremely diverse in regulation of cellular communication, which is fundamental to all organisms and mediates numerous processes, such as cell fate decisions, proliferation, migration, and homeostasis [31]. In insects, a subseries of pathways was identified to date, including Notch, Wnt, Hedgehog, TGF-β, Hippo, NF-*κ*B, and JAK/STAT signaling pathways [31]. TGF-β signaling pathway is highly correlated with expression of ncRNAs such as miRNAs and lncRNAs, TGF-β indirectly represses miR-200 family members through induction of ZEB1 and ZEB2 transcription factors, and miR-200 family members target and inhibit the expression of ZEB proteins, which stabilize regulatory networks for TGF-β-induced epithelial-mesenchymal transition [32-33].In the present study, we observed that TGF-β signaling pathway-related mRNAs were targeted by four DEpiRNAs (piR-bmo-748815, piR-bmo-1183555, piR-bmo-458012, and piR-bmo-512574) in the Ac4 vs. Ac5 comparison group and 18 DEpiRNA (piR-bmo-1188284, piR-bmo-934193, and piR-bmo-1183075, etc.) in the Ac5 vs. Ac6 comparison group. This indicated that the aforementioned DEpiRNAs were likely to participate in controlling TGF-β signaling pathway by modulating downstream gene expression, further regulating the larval gut development.

As an important downstream mediator for a variety of eukaryotes, Jak-STAT signaling pathway plays an essential role in metabolism. Dodington et al. found that JAK-STAT signaling in the peripheral metabolic organs has been shown to regulate a multitude of metabolic processes with the use of tissue-specific knock-out mice [34]. In *Drosophila* gut, JAK/STAT signaling is essential for intestinal stem cell differentiation and for intestinal regeneration after insults and infection [35]. Here, we found Jak-STAT signaling pathway-related genes were targeted by five DEpiRNAs (piR-bmo-748815, piR-bmo-1183555, piR-bmo-1103846, piR-bmo-512574 and piR-bmo-202265) in the Ac4 vs. Ac5 comparison group and 18 DEpiRNA (piR-bmo-1188284, piR-bmo-934193 and piR-bmo-1183075, etc.) in the Ac5 vs. Ac6 comparison group. This indicated that the aforementioned DEpiRNAs have high correlation with Jak-STAT signaling pathway, which were likely to participate in metabolism process of midgut during *A. c. cerana* larvae development.

Previously, overexpression and knockdown of piRNAs by feeding or injection were verified to be effective in several animals such as *Culex pipiens pallens, Nematostella vectensis*., and *Bemisia tabaci* [36-38]. In the near future, we will conduct functional investigation of candidate DEpiRNAs identified in this work through overexpression and knockdown.

## 5. Conclusion

Taken together, 621 piRNAs were for the first time identified in the *A. c. cerana* larval guts. The overall expression pattern of piRNAs changed during the developmental process of larval guts were identified. DEpiRNAs potentially modulate the development of *A. c. cerana* larval guts by regulating an array of functional terms and pathways, such as Wnt, TGF-β, and Jak-STAT signaling pathways.

## Supplementary Materials

The following supporting information can be downloaded at: www.mdpi.com/xxx/s1, Table S1: Primers that were used in this study; Table S2: Detailed information about the identified piRNA in *A. c. cerana* larvae; Table S3: Table S3. Detailed information about the targeting relationship between DEpiRNAs and mRNAs; Figure S1: Ridgeline plots of total piRNAs expression level in *A. c. cerana* and *A. m. ligustica*

## Author Contributions

R.G. and D.C. designed this research; Q.L., M.S. contributed to the writing of the article; Q.L., M.S., X.F., W.Z., D.Z., Y.H., Z.W., K.Z., K.Y., H.Z., Y.S. and Z.F., conducted experiments and data analyses. R.G., D.C and Q.N. supervised the study and preparation of the manuscript.

## Acknowledgments

We thank all editors and reviewers for their helpful and constructive comments. RG appreciates his adorable daughter for her love and goodness.

